# Tracking insulin- and glucagon-expressing bihormonal cells during differentiation using an *INSULIN* and *GLUCAGON* double reporter human embryonic stem cell line

**DOI:** 10.1101/2023.04.19.537542

**Authors:** Samantha Mar, Ekaterina Filatov, Cuilan Nian, Shugo Sasaki, Dahai Zhang, Francis C. Lynn

## Abstract

Human embryonic stem cell (hESC)-derived pancreatic alpha and beta cells can be used to develop cell replacement therapies to treat diabetes. However, recent published differentiation protocols yield varying amounts of alpha and beta cells amidst heterogeneous cell populations. To visualize and isolate hESC-derived alpha and beta cells, we generated a *GLUCAGON-2A- mScarlet* and *INSULIN-2A-EGFP* dual fluorescent reporter (INS^EGFP^GCG^mScarlet^) hESC line using CRISPR/Cas9. We established robust expression of EGFP and mScarlet fluorescent proteins in insulin- and glucagon-expressing cells respectively without compromising the differentiation or function of these cells. We also showed the insulin- and glucagon-expressing bihormonal population at the maturing endocrine cell stage (Stage 6) lose insulin expression over time, while maintaining an alpha-like expression profile, suggesting these bihormonal cells are preferentially fated to become alpha-like cells *in vitro*. Together, the INS^EGFP^GCG^mScarlet^ hESC line provides an efficient strategy for tracking populations of hESC-derived beta- and alpha-like cells.

## Introduction

Pancreatic beta and alpha cells secrete hormones insulin and glucagon respectively to regulate glucose homeostasis in the blood. These cells organize into clusters during pancreas development, forming the islets of Langerhans. People living with type 1 diabetes (T1D) lack sufficient beta cells, resulting in elevated blood glucose levels that can lead to life-threatening complications. Thus, it is critical for these people to rely on exogenous insulin to manage their disease long-term. Transplantation of cadaveric human islets have been shown to effectively restore regulated insulin production in people living with T1D, allowing patients to become independent of exogenous insulin (Shapiro, 2012; Shapiro et al., 2017). However, there is a scarce supply of donor islets and the quality of these islets is variable, limiting the accessibility of this treatment (Bellin et al., 2012; Sneddon et al., 2018). Therefore, human embryonic stem cells (hESCs) have been proposed to serve as an unlimited and reliable source of pancreatic beta and alpha cells for the treatment of diabetes (Shapiro & Verhoeff, 2023).

hESC lines engineered to report insulin using a fluorescent protein have proved to be an efficient tool used to further improve the generation of pancreatic beta cells *in vitro* (Micallef et al., 2012; Nair et al., 2019; Novakovsky et al., 2023). These fluorescence protein-expressing cells can easily be visualized using a benchtop fluorescence microscope, used in high throughput screens or isolated using fluorescence activated cell sorting (FACS) for downstream analyses (Giudice & Trounson, 2008). However, recently published differentiation protocols generate heterogeneous cultures, with varying amounts of insulin-expressing cells, glucagon-expressing cells, as well as insulin- and glucagon-expressing bihormonal cells (Balboa et al., 2022; Nair et al., 2019; Russ et al., 2015; Velazco-Cruz et al., 2019). Thus, an insulin single reporter hESC line would not be able to differentiate between these monohormonal and bihormonal cells.

To visualize and isolate monohormonal and bihormonal cell populations during differentiation, we generated an insulin and glucagon dual fluorescent reporter hESC line using CRISPR/Cas9. Specifically, we knocked-in added on *2A-EGFP* and *2A-mScarlet* expression cassettes downstream of the *INS* and *GCG* loci of the H1/WA01 hESC line. To validate the cell line, we showed EGFP and mScarlet expression report insulin and glucagon expression respectively. We confirmed the differentiation and function of these reporter-derived cells were not affected by the insertion of the expression cassettes. To demonstrate this cell line can be used to enrich beta- and alpha-like cells, we sorted cells based on fluorescence reporter expression and found distinct populations that expressed key beta and alpha cell markers. Interestingly, we also found the expression profile of the insulin- and glucagon-expressing cell population at the maturing endocrine cell stage more closely resembles that of alpha cells than beta cells. To test whether this population will further differentiate into monohormonal, glucagon-expressing alpha cells, we reaggregated these bihormonal cells for further culture and found that the majority of these cells lose insulin expression over time. These findings suggest these bihormonal cells represent a transient population that are preferentially fated to become monohormonal alpha-like cells.

## Materials and Methods

### Cell Culture

Stem cells were maintained as previously described (Huang et al., 2023). Briefly, undifferentiated H1 (WA01; XY) hESCs were maintained in mTeSR Plus (STEMCELL Technologies Inc.) at 5% CO_2_, 37°C. Cells were split every 4 days using ReLeSR (STEMCELL Technologies Inc.) and plated 1×10^6^ and 2×10^6^ cells on Geltrex (Gibco, 1:100 DMEM/F12, Thermo Fisher Scientific)- coated 60 mm and 10 cm plates respectively, with 10 μM Y-27632 (STEMCELL Technologies Inc.).

### CRISPR/Cas9 Knock-In

The CRISPR/Cas9 system used to generate the INS^EGFP^GCG^mScarlet^ hESC line was as previously described (Huang et al., 2023; Krentz et al., 2014; Novakovsky et al., 2023). The pCCC CRISPR/Cas9 plasmids were generated to express gRNA (5’-TGCAACTAGACGCAG- 3’) and (5’-AACATCACCTGCTAGCCACG-3’) for targeting the 3’ end of the endogenous insulin (*INS*) and glucagon (*GCG*) coding regions respectively. The homology arms of the selected CRISPR clones were genotyped and subsequently sequenced to confirm fidelity for recombination (Figure S1A). For a list of genotyping primers used, see Table S1. Clones with the insertions were then tested for common chromosomal abnormalities using the hPSC Genetic Analysis Kit (STEMCELL Technologies Inc.).

### hESC Differentiation

The protocol used to differentiate the hESC lines was based on previously published protocols (Balboa et al., 2022; Huang et al., 2023; Nair et al., 2019; Russ et al., 2015). Briefly, hESCs were dissociated using Accutase (STEMCELL Technologies Inc.) for 5 minutes at 37°C and washed using DPBS with calcium and magnesium (Gibco). One million cells/mL was seeded in a total of 5.5 mL mTesr Plus containing 10 μM Y-27632 in each well of non-tissue culture treated 6-well plates (Greiner Bio-One). The plates were placed on a shaker (25 mm orbit; Celtron, Infors HT) set at 94 RPM at 37°C, 5% CO_2_ overnight. The 3D spheroids were washed using DPBS with calcium and magnesium and replaced with differentiation media. For detailed formulations of differentiation media at each stage, see Table S2.

### Flow Cytometry and FACS

For flow cytometry analysis, spheroids were washed with PBS and dissociated using Accutase (STEMCELL Technologies Inc.) for 8 minutes at 37°C. Dissociated cells were centrifuged for 5 minutes at 200 x g and resuspended in Fixable Viability Dye eFluor 780 (Thermo Fisher Scientific, 1:1000 in PBS) for 30 minutes at 22°C. Cells were subsequently centrifuged for 5 minutes at 200 x g and resuspended in 4% paraformaldehyde (Thermo Scientific Chemicals) for 30 minutes at 22°C. Cells were then centrifuged for 5 minutes at 200 x g and resuspended in PBS for long term storage. On the day of analysis, cells were permeabilized using 0.5% triton X-100 (Thermo Fisher Scientific) and stained with conjugated antibodies for 1 hour on ice: mouse anti-C-peptide antibody conjugated with Alexa Fluor 647 (BD Biosciences) and mouse anti-glucagon antibody conjugated with Bv421 (BD Biosciences). Cells were analyzed on a BD LSR II using unstained and single stain controls, and the data was processed using FlowJo. To sort mScarlet^-^EGFP^lo^, mScarlet^-^EGFP^hi^, mScarlet^+^EGFP^-^, and mScarlet^+^EGFP^hi^ populations, spheroids were rinsed with PBS and dissociated using Accumax for 8 minutes at 37°C. Dissociated cells were resuspended in 2% FBS (HyClone, Thermo Fisher Scientific) in PBS with 10 μM Y-27632 on ice and sorted into Stage 7 media with 10 μM Y- 27632 using the BD FACS Aria.

### Gene Expression Analyses

RNA extraction and RT-qPCR using sorted cells was performed as previously described (Sabatini et al., 2018). ΔΔCT was normalized to the housekeeping gene *TBP*. For the list of TAQMAN primers used, see Table S3. NanoString gene expression analysis was performed as previously described (Huang et al., 2023). 50,000 sorted cells were resuspended in 100 μL Buffer RLT (Qiagen) containing 1% β-mercaptoethanol (Sigma-Aldrich). The prepared sample cartridge was read on a NanoString nCounter SPRINT profiler and analyzed using nSolver 4.0 for relative gene expression. Data was normalized to six housekeeping genes: *B2M*, *GAPDH*, *GUSB*, *HPRT1*, *POLR2A*, and *TBP*. For the list of target gene sequences, see Table S4.

### Immunohistochemical Analyses

Briefly, 100-200 spheroids were fixed in 4% PFA for 30 minutes at 22°C, embedded in 2% agarose. The agarose containing the spheroids was then dehydrated and embedded into paraffin blocks. The paraffin blocks were sectioned into 5 μm sections. The sections were then rehydrated, heated in antigen retrieval buffer (0.433% v/v citraconic anhydride 98% in dH_2_O, Alfa Aesar), and blocked for 30 minutes in 5% v/v horse serum in PBS. The sections were stained with primary antibodies diluted in 5% v/v horse serum over night at 4°C: mouse anti-glucagon antibody (1:1000, Sigma-Aldrich), guinea pig anti-insulin (1:500, Agilent), rat anti-RFP (1:1000, ChromoTek), and rabbit anti-GFP (1:500, MBL). Then, the sections were stained with secondary antibody for 1 hour at 22°C in the dark: anti-mouse Cy3 (1:250, Jackson ImmunoResearch), anti-guinea pig Alexa Fluor 594 (1:250, Jackson ImmunoResearch), anti-rat Rhodamine Red X (1:250, Jackson ImmunoResearch), anti-rabbit FITC (1:450, Jackson ImmunoResearch), and TO-PRO-3 (1:10,000, Thermo Fisher Scientific). Images were taken using a 20x oil immersion and a 10x dry objective on a Leica TCS SP8 confocal microscope.

### Hormone Secretion Assay

Fifty size-matched spheroids were used per treatment and pre-incubated in 500 μL Krebs-ringer bicarbonate HEPES supplemented with 2.8 mM D-glucose or 5 mM D-glucose (Sigma-Aldrich) for 1 hour for glucose-stimulated insulin secretion and glucose-mediated glucagon secretion assays respectively. After preincubation, the spheroids were transferred into wells containing 250 μL of the treatment solution and incubated for 1 hour at 5% CO_2_, 37°C. For glucose-stimulated insulin secretion, the treatments solutions were 250 μL Krebs-ringer bicarbonate HEPES that contained 2.8 mM D-glucose, 16 mM D-glucose, or 16 mM D-glucose with 30 mM KCl. For glucagon secretion, the treatment solutions contained 5 mM D-glucose, 1 mM D-glucose, or 1 mM D-glucose with 30 mM Arginine. After the incubation, the supernatants were centrifuged at 5,000 x g for 10 minutes and collected. The spheroids were transferred into 500 μL acid ethanol (1 M HCl in 70% ethanol) for either human C-peptide ELISA (Alpco) or human glucagon ELISA (Mercodia).

### Statistical Analyses

Statistical analyses were performed using Prism 9 (GraphPad Software). All data are presented as mean ± s.e.m. Data were analyzed using either a Student’s *t* test or a one-way ANOVA followed by a Dunnett post-hoc test for multiple comparisons where appropriate. Significance was determined using p < 0.05.

## Results

### Generation of the INS^EGFP^GCG^mScarlet^ hESC reporter line using CRISPR/Cas9

We generated a reporter hESC line using CRISPR/Cas9-mediated knock-in to identify and isolate insulin- and glucagon-expressing cells in hESC-derived pancreatic endocrine cell cultures. We first developed the single knock-in INS^EGFP^ hESC line by inserting a *2A-EGFP* expression cassette into exon 3 of the endogenous *INS* gene in the H1 hESC line (Figure 1A) (Novakovsky et al., 2023). We specifically replaced the stop codon of the *INS* gene to prevent disruptions to insulin processing in *INS*-expressing cells (Blöchinger et al., 2020). The *2A* sequence that promotes ribosomal skipping was used to generate a bi-cistronic reporter system that allows equimolar protein expression of INS and EGFP to further minimize any disruptions to insulin processing (Blöchinger et al., 2020; Liu et al., 2017). The double knock-in INS^EGFP^GCG^mScarlet^ hESC line was generated using the INS^EGFP^ hESC line in a similar manner. We introduced a *2A-mScarlet* expression cassette downstream of the glucagon (*GCG*) gene using the INS^EGFP^ Clone #26 hESC line, replacing the STOP codon of the endogenous *GCG* gene (Figure 1B).

**Figure 1:**
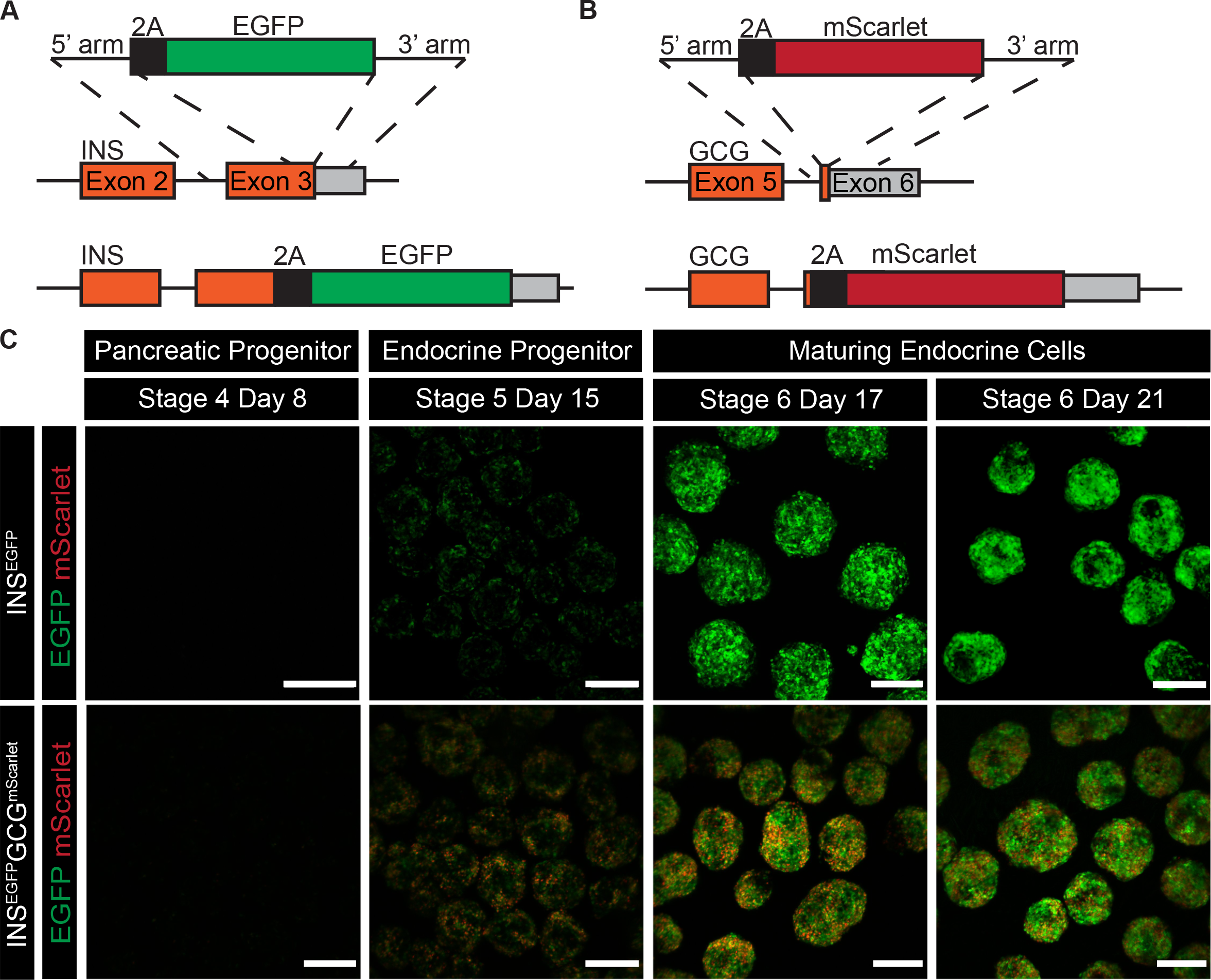
Generation of INS^EGFP^ and INS^EGFP^GCG^mScarlet^ reporter human embryonic stem cell (hESC) lines using CRISPR-Cas9. (A,B) Schematic overview of the CRISPR-Cas9 targeting strategy to knock-in (A) a *2A-EGFP* expression cassette onto exon 3 of the *INS* gene and (B) a *2A-mScarlet* expression cassette onto exon 6 of the *GCG* gene of the wildtype H1 hESC line. (C) Differentiated spheroids derived from the INS^EGFP^ and INS^EGFP^GCG^mScarlet^ reporter cell lines expressing EGFP (green) and mScarlet (red) at different stages of pancreatic endocrine cell differentiation. Images were obtained on a Leica SP8 confocal microscope. Scale bars = 200 *μ*m.

### INS^EGFP^GCG^mScarlet^ reporter line reports INS and GCG in hESC-derived pancreatic endocrine cells without affecting their differentiation capacity

To ensure the genomic modifications do not affect the differentiation of hESC-derived pancreatic endocrine cells, we followed the differentiation of pancreatic endocrine cells derived from the single and double knock-in hESC lines. We observed the morphology of the spheroids derived from both genetically modified cell lines remained comparable to those generated using previously published differentiation protocols (Figure S1A,B,C). We also found EGFP and mScarlet reporter are expressed when *INS* and *GCG* are expected to be expressed, namely starting at the endocrine progenitor cell stage (Stage 5), and persisting throughout the maturation process of pancreatic endocrine cells (Figure 1C). Of our total culture, we found 2.55% ± 0.79% express mScarlet only (mScarlet^+^), 23.90% ± 3.49% express both EGFP and mScarlet (EGFP^+^mScarlet^+^), and 35.12% ± 3.81% express EGFP only (EGFP^+^) at the maturing endocrine cell stage (Stage 6 Day 25) using the INS^EGFP^GCG^mScarlet^ Clone E7, which is comparable to the differentiation efficiencies of two other INS^EGFP^GCG^mScarlet^ clones (Figure S2A). These data suggest the insertion of the expression cassettes does not affect the morphology, hormone expression or differentiation efficiency of the hESC-derived pancreatic endocrine cell clusters at each stage of the differentiation.

To determine if the fluorescent reporter proteins report INS and GCG expression, maturing endocrine cells derived from the knock-in hESC lines were immunostained for these hormones and their corresponding fluorescent proteins. Using immunofluorescence microscopy, we saw insulin and EGFP co-expression in the single and double knock-in line, indicating EGFP reports insulin hormone expression (Figure 2A). Flow cytometry analysis also showed 96.54% ± 1.12% and 97.7% ± 0.43% C-peptide-expressing cells also express EGFP for the INS^EGFP^ and INS^EGFP^GCG^mScarlet^ cell lines respectively at the maturing endocrine cell stage (Stage 6 Day 21) (Figure 2C and Figure S2B,C). Similarly, we observed glucagon and mScarlet co-expression in the double knock-in line and the mScarlet reporter efficiency was 83.94% ± 4.76%, demonstrating mScarlet reports glucagon hormone expression (Figure 2B,D and Figure S2D). Together, EGFP and mScarlet reports expression of insulin and glucagon in these knock-in cell lines.

**Figure 2:**
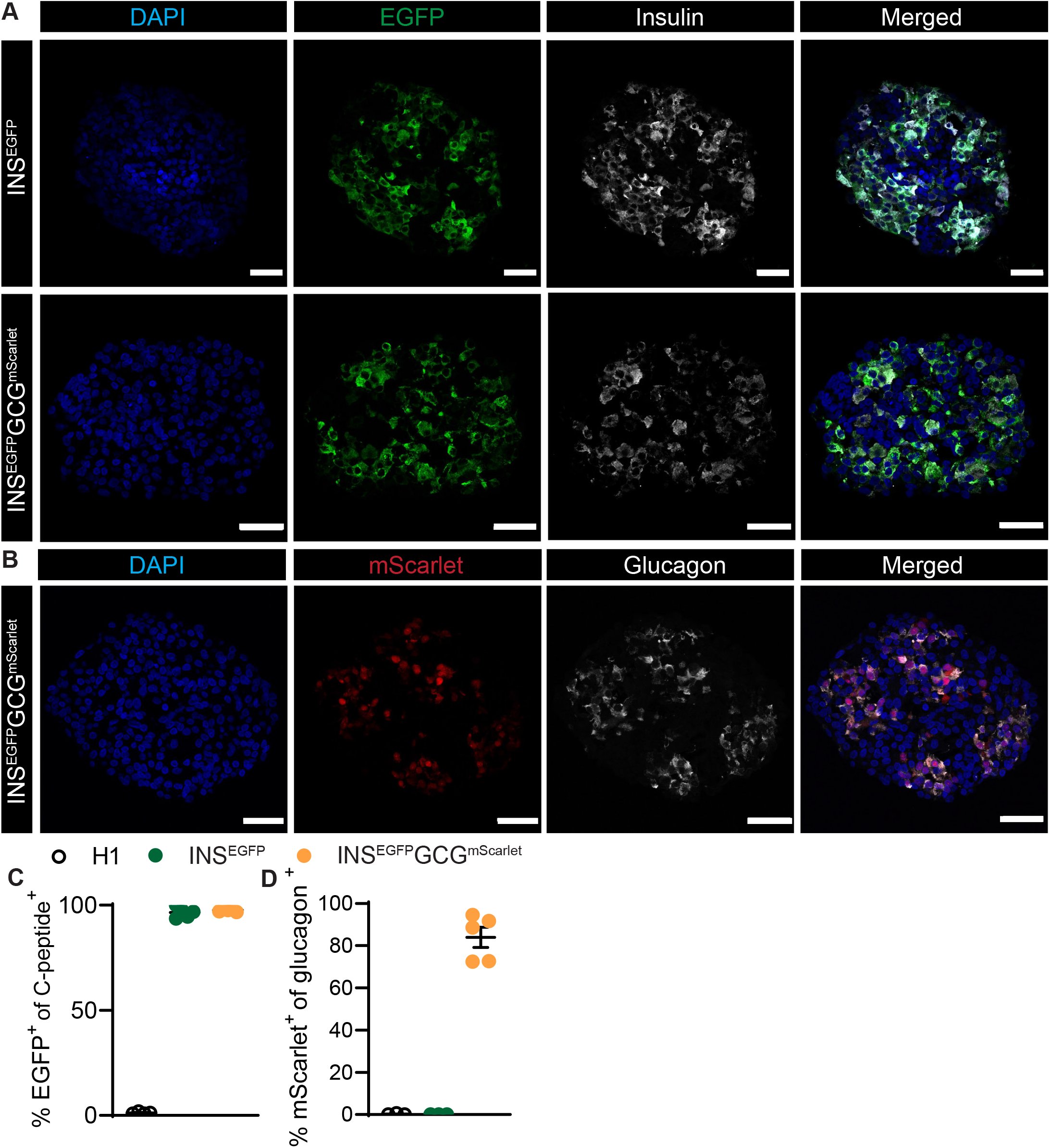
EGFP and mScarlet expression report insulin and glucagon hormone expression. (A) Immunostaining of EGFP (green), insulin (grey), and nuclei (blue) of maturing endocrine cells (Stage 6 Day 21) derived from the INS^EGFP^ and INS^EGFP^GCG^mScarlet^ reporter cell lines. n = 3, scale bars = 50 *μ*m. (B) Immunostaining of mScarlet (red), glucagon (grey), and nuclei (blue) of maturing endocrine cells (Stage 6 Day 21) derived from the INS^EGFP^GCG^mScarlet^ reporter cell line. n = 3, scale bars = 50 *μ*m. (C,D) Quantitative flow cytometry analysis of the reporter efficiency of maturing endocrine cells (Stage 6 Day 21) derived from the H1 parental, INS^EGFP^ and INS^EGFP^GCG^mScarlet^ reporter cell lines, measuring percentage of (C) EGFP^+^ cells of C-peptide^+^ cells, and (D) mScarlet^+^ cells of glucagon^+^ cells. n = 3-5, error bars report ±SEM.

### INS^EGFP^GCG^mScarlet^ reporter line can enrich for beta- and alpha-like cell populations

To determine if the fluorescent reporter proteins can be used to effectively isolate beta- and alpha-like cell populations, we sorted distinct fluorescence cell populations using FACS for gene expression analysis using NanoString (Figure 3A). We noticed there were two distinct EGFP-expressing populations, so we further split the EGFP^+^ population into two groups, cells with high (EGFP^hi^) and low (EGFP^lo^) fluorescence intensities. For comparison purposes, we gated the EGFP and mScarlet double positive population to correspond to that of EGFP^hi^ (EGFP^hi^mScarlet^+^). We also defined the gating of our populations more stringently as to enrich for more pure cell populations. Our NanoString analysis showed the transcriptomic profile of EGFP^hi^mScarlet^+^ and EGFP^hi^ populations more closely resemble primary beta and alpha cells within our cultures as they express higher levels of the endocrine hormones *INS* and *GCG*, the pan-endocrine marker *CHGA*, and the hormone secretion machinery *SLC30A8* compared to the two other populations. We also found the EGFP^hi^mScarlet^+^ cells adopt a more alpha-like expression profile with higher expression levels of key alpha cell regulator *ARX* and lower levels of key beta cell regulator *NKX6-1* compared to the EGFP^hi^ population, despite expressing similar levels of *INS* (Figure 3B-E). On the other hand, the EGFP^hi^ population is more beta-like as it expresses key beta cell regulators such as *NKX6-1*, *PCSK1*, and *PAX4*, at significantly higher levels than the EGFP^hi^mScarlet^+^ population (Figure 3D, F, H).

**Figure 3:**
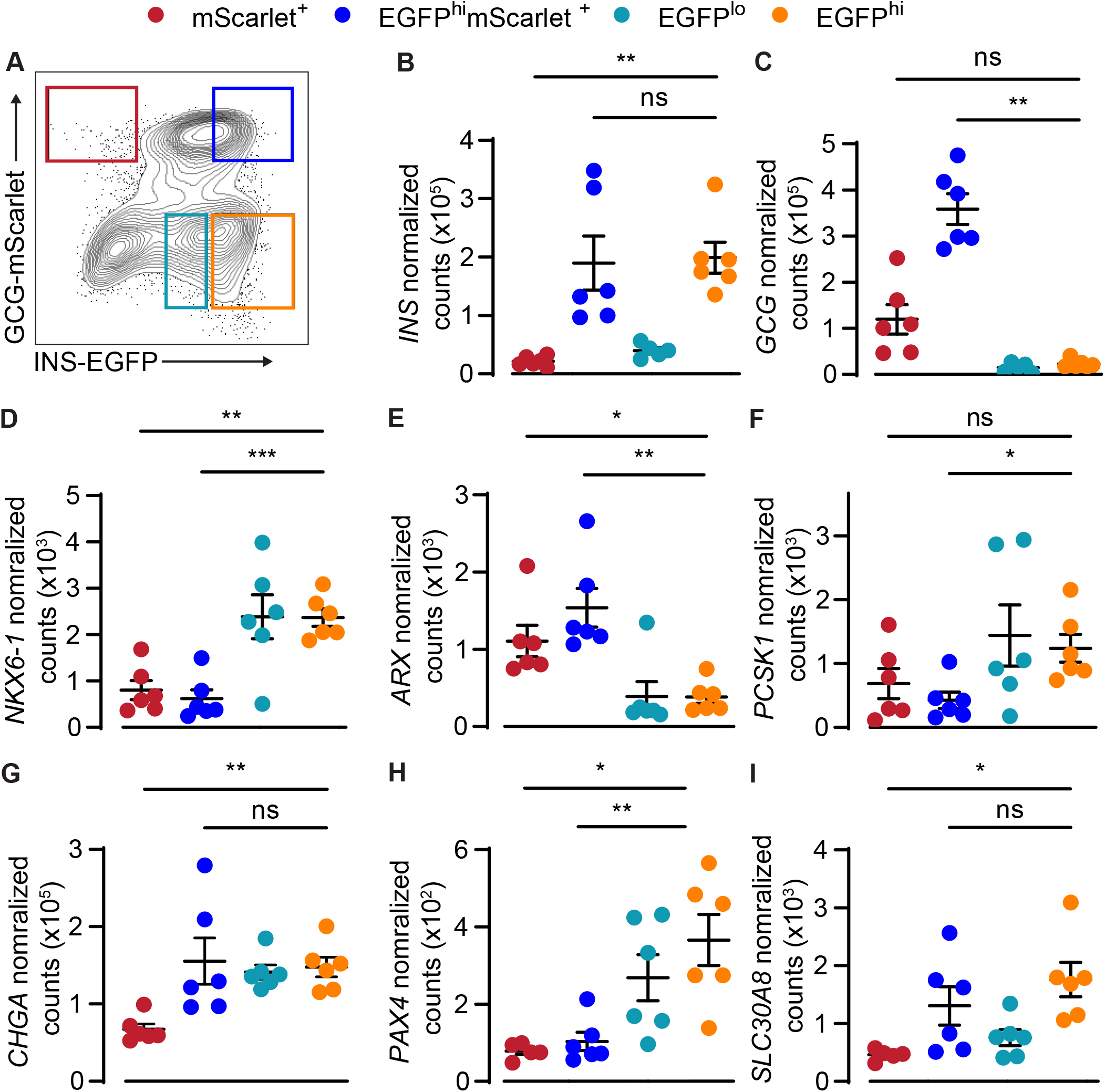
INS^EGFP^GCG^mScarlet^ derived maturing endocrine cells can be used to enrich populations with distinct transcriptional profiles using fluorescence-activated cell sorting (FACS). (A) FACS plot of maturing pancreatic endocrine cells derived from the INS^EGFP^GCG^mScarlet^ reporter cell line (Stage 6 Day 25) (B-I) Transcript expression of (B) *INS*, (C) *GCG*, (D) *NKX6-1*, (E) *ARX*, (F) *PCSK1*, (G) *CHGA*, (H) *PAX4*, and (I) *SLC30A8* in FACS-sorted maturing pancreatic endocrine cells (Stage 6 Day 25). n = 6, * = *p* ≤ 0.0332, ** = *p* ≤ 0.0021, one-way ANOVA, error bars report ±SEM.

To determine if EGFP^hi^mScarlet^+^ cells are fated to become alpha-like cells, this cell population was sorted and reaggregated at the maturing endocrine cell stage (Figure 4A). Confocal imaging and flow cytometry analysis showed there were significantly less EGFP^hi^mScarlet^+^ cells over time in unsorted differentiation cultures (Figure 4B and 5B). We also found the sorted and reaggregated EGFP^hi^mScarlet^+^ cell population became mostly mScarlet^+^, while the sorted and reaggregated EGFP^hi^ population remained mostly EGFP^+^ (Figure 4B, C). RT-PCR analysis further demonstrated the sorted and reaggregated EGFP^hi^mScarlet^+^ cells expressed lower levels of *INS* and higher levels of *GCG*, *ARX*, and *IRX2* compared to the EGFP^hi^ cells, suggesting EGFP^hi^mScarlet^+^ cells are preferentially fated to become alpha-like cells (Figure 4D).

**Figure 4:**
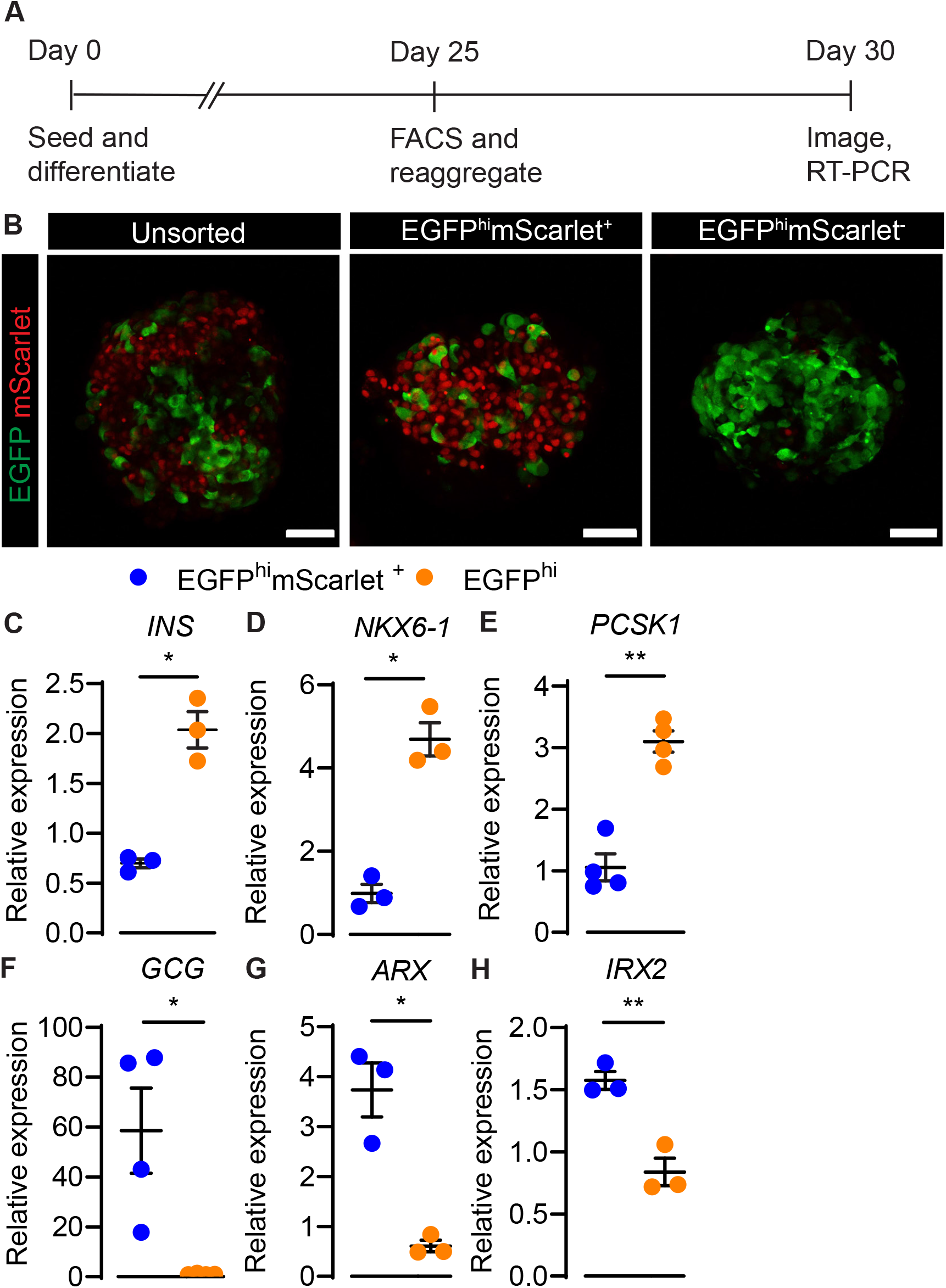
EGFP^hi^mScarlet^+^ sorted and reaggregated cells are fated to become alpha-like cells. (A) Schematic of experimental methodology. (B) Unsorted, and sorted and reaggregated EGFP^hi^mScarlet^+^ and EGFP^hi^ populations 5-days post-sort. n = 3, scale bars = 50 *μ*m (C-H) Transcript analysis of (C) *INS*, (D) *NKX6-1*, (E) *PCSK1*, (F) *GCG*, (G) *ARX*, and (H) *IRX2* of FACS-sorted and reaggregated maturing endocrine cells derived from INS^EGFP^GCG^mScarlet^ reporter cell line. n = 3-4, * = *p* ≤ 0.0332, ** = *p* ≤ 0.0021, paired t-test, error bars report ±SEM.

**Figure 5:**
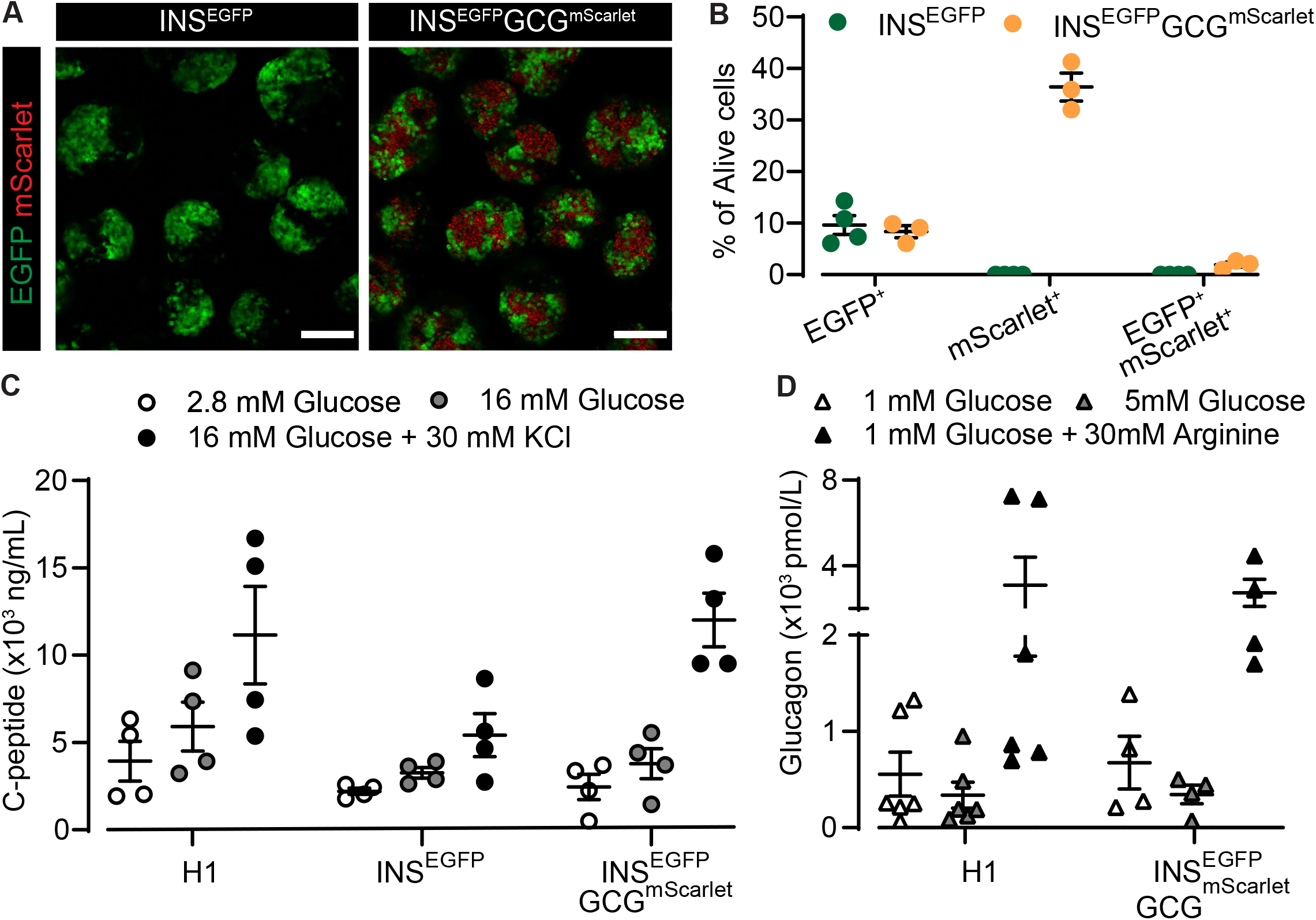
Targeting the *INS* and *GCG* loci does not affect the function of hESC-derived pancreatic β- and a-like cells. (A) Mature pancreatic endocrine cells (Stage 7 Day 44-45) derived from the INS^EGFP^ and INS^EGFP^GCG^mScarlet^ reporter cell lines. Images were obtained on a Leica SP8 confocal microscope. Scale bars = 200 *μ*m. (B) Differentiation efficiency of mature pancreatic endocrine cells (Stage 7 Day 45-55) derived from INS^EGFP^ and INS^EGFP^GCG^mScarlet^ reporter cell lines determined using flow cytometry analysis. n = 3-4, error bars report ±SEM. (C) Glucose-stimulated insulin secretion assay using mature pancreatic endocrine cells (Stage 7 Day 50-60) derived from the H1 parental, INS^EGFP^, and INS^EGFP^GCG^mScarlet^ reporter cell lines. n = 4, error bars report ±SEM. (D) Glucose-mediated glucagon secretion assay using mature pancreatic endocrine cells (Stage 7 Day 45-55) derived from the H1 parental and INS^EGFP^GCG^mScarlet^ reporter cell lines. n = 4-6, error bars report ±SEM.

### INS^EGFP^GCG^mScarlet^ reporter line can differentiate into functionally mature pancreatic endocrine cells

To determine whether the genetic modifications affect the functional maturation of hESC-derived pancreatic endocrine cells, we differentiated the INS^EGFP^ and INS^EGFP^GCG^mScarlet^ cell lines into mature pancreatic endocrine cells (Figure 5A). At this stage, we observed 9.63% ± 1.85% and 8.35% ± 1.17% express EGFP only using INS^EGFP^ and INS^EGFP^GCG^mScarlet^ cell lines respectively, and 36.4% ± 2.70% express mScarlet only using the INS^EGFP^GCG^mScarlet^ cell line (Figure 5B). To test whether these cells are functionally mature, we performed glucose-stimulated insulin and glucagon secretion tests on mature pancreatic endocrine cells generated using the reporter hESC lines and compared the results to cells generated using the H1 parental cell line. We found C-peptide secretion was stimulated at high glucose concentrations using all three cell lines (Figure 5C). We also observed these cultures secrete less glucagon at normoglycemic glucose levels compared to that of low glucose levels, and the glucagon secretion levels are comparable between cell lines (Figure 5D). These data suggest the genetic modifications do not affect hormone secretion and maturation of hESC-derived pancreatic endocrine cells.

## Discussion

Current pancreatic endocrine cell differentiation protocols generate heterogeneous cultures. Here we describe the generation and characterization of the INS^EGFP^GCG^mScarlet^ reporter hESC line for the visualization and enrichment of hESC-derived beta- and alpha-like cells. We validated this double knock-in reporter line reports insulin and glucagon expression with EGFP and mScarlet expression. Our phenotypic and functional studies of the cell line also show that the genetic modifications do not disrupt endogenous insulin and glucagon expression or secretion. We also demonstrated that this reporter line can enrich alpha- and beta-like cells as well as track their differentiation. Thus, the INS^EGFP^GCG^mScarlet^ hESC reporter line can be used to study beta and alpha cells during pancreatic endocrine cell differentiations and further improve the production of functional hESC-derived pancreatic endocrine cells for diabetes therapy.

Insulin- and glucagon-expressing bihormonal cells have previously been described in the developing human pancreas (Riedel et al., 2012; Riopel et al., 2014). These cells were shown to become less frequently observed over time in the human fetal pancreas, and their numbers become negligible in adulthood (Riedel et al., 2012; Riopel et al., 2014). Based on the transcription factor profile of these bihormonal cells, it has been suggested that these cells are alpha cell precursors (Riedel et al., 2012). Similar observations have been made in hESC-derived polyhormonal cells. Differentiation protocols generated by other groups found a similar reduction in the number of insulin- and glucagon-expressing bihormonal cells accompanied by an increase in the number of glucagon-expressing monohormonal cells (Balboa et al., 2022; Peterson et al., 2020; Rezania et al., 2011; Veres et al., 2019). Transcriptomic analyses have also shown these bihormonal cells possess a more alpha-like expression profile (Augsornworawat et al., 2020; Balboa et al., 2022; Basford et al., 2012; Peterson et al., 2020; Veres et al., 2019). To determine whether these bihormonal cells further differentiate into monohormonal alpha-like cells, others have sorted and transplanted polyhormonal cells into mice, and found these cells eventually resolve into monohormonal alpha-like cells (Alvarez-Dominguez et al., 2020; Basford et al., 2012; Kelly et al., 2011; Rezania et al., 2013). However, previous sorting strategies used to enrich polyhormonal cells for transplantation do not yield a pure insulin- and glucagon- expressing bihormonal cell population so it is unclear whether these bihormonal cells are fated to become alpha-like cells. Using the INS^EGFP^GCG^mScarlet^ reporter line, we showed that these *in vitro* generated bihormonal cells lose insulin expression and maintain an alpha-like transcriptional profile during maturation *in vitro*. The dual reporter line can be used to further examine these cell fate decisions *in vitro* and *in vivo*.

Human islets commonly adopt a mixed cytoarchitecture where alpha and beta cells intermingle. Previous studies have shown dispersed human islets do not self-organize into their native islet architecture, but instead adopt a core-mantle architecture, reducing the cell-cell interactions between alpha and beta cells (Lavallard et al., 2016). This observation suggests properties intrinsic to human islet cells are not enough to recapitulate their *in vivo* architecture (Adams & Blum, n.d.; Lavallard et al., 2016). At the mature endocrine cell stage of our differentiation protocol, alpha-like and beta-like cells are clustered separately with very little cell-cell contact between the two cell types, similar to dispersed human islets. Thus, the dual reporter line could be used to determine extrinsic factors that contribute to human islet architecture and these factors may be targeted to better mimic *in vivo* architecture *in vitro*.

In summary, we generated an INS^EGFP^GCG^mScarlet^ reporter hESC line that reports insulin and glucagon without affecting the differentiation or function of hESC-derived beta- and alpha-like cells. We also showed that insulin- and glucagon-expressing cells can be tracked and purified using the INS^EGFP^GCG^mScarlet^ hESC reporter line. This dual reporter line could therefore be used to further characterize and improve *in vitro* pancreatic endocrine differentiations for diabetes therapy.

## Acknowledgements

The authors thank the members of the Lynn, Levings, Verchere, Wasserman, Johnson and Kieffer Laboratories (Vancouver, British Columbia, Canada); the Bruin Laboratory (Carleton University); the PE MacDonald Laboratory (University of Alberta); and the BCCHR Flow Cytometry and Imaging core facilities for technical support, discussion, and critical reading of the manuscript. F.C.L. was supported by the JDRF (5-SRA-2020-1059-S-B, 3-COE-2022-1103- M-B) and Canadian Institutes of Health Research (ASD-173663). Salary (F.C.L.) was supported by the Michael Smith Foundation for Health Research (#5238 BIOM) and the BC Children’s Hospital Research Institute. Scholarship funding was provided by the Canadian Institutes of Health Research (CGSM; S.M., E.F.), The BC Children’s Hospital Research Institute (S.M., E.F.), the UBC CELL Graduate Program (1YF; S.M.) and the Canadian Islet Research and Training Network NSERC CREATE program (E.F.). The authors acknowledge that UBC and BC Children’s Hospital are situated on the traditional, ancestral, and unceded territories of the Coast Salish peoples, the Sḵwx̱ wú7mesh (Squamish), sə^l̓^ ilwətaɁɬ (Tsleil-Waututh), and xʷməθkʷə^y̓^ əm (Musqueam) Nations.

---

**Figure S1:**
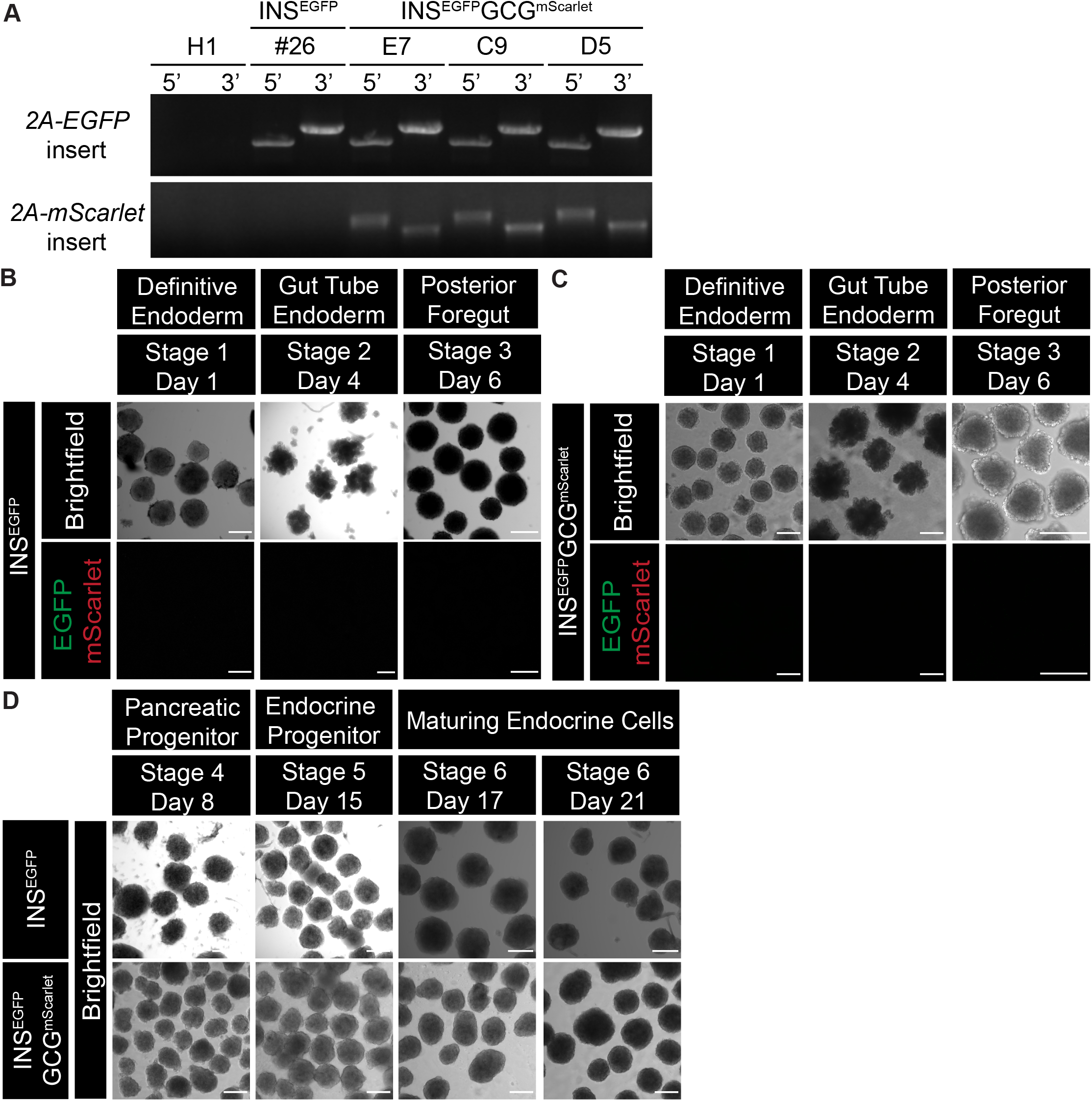
Generation of INS^EGFP^ and INS^EGFP^GCG^mScarlet^ reporter human embryonic stem cell (hESC) lines using CRISPR/Cas9. (A) Genotyping PCR for 5’ and 3’ homology arm of the *2A-EGFP* and *2A-mScarlet* insertion for wildtype H1 parental, INS^EGFP^ clone #26, and three of the INS^EGFP^GCG^mScarlet^ clones E7, C9 and D5. (B,C) Differentiated spheroids derived from the (B) INS^EGFP^ and (C) INS^EGFP^GCG^mScarlet^ reporter cell lines expressing EGFP (green) and mScarlet (red) at different stages of pancreatic endocrine cell differentiation. Scale bars = 200 *μ*m. (D) Brightfield images of differentiated spheroids derived from the INS^EGFP^ and INS^EGFP^GCG^mScarlet^ reporter cell lines at different stages of pancreatic endocrine cell differentiation. Scale bars = 200 *μ*m.

**Figure S2:**
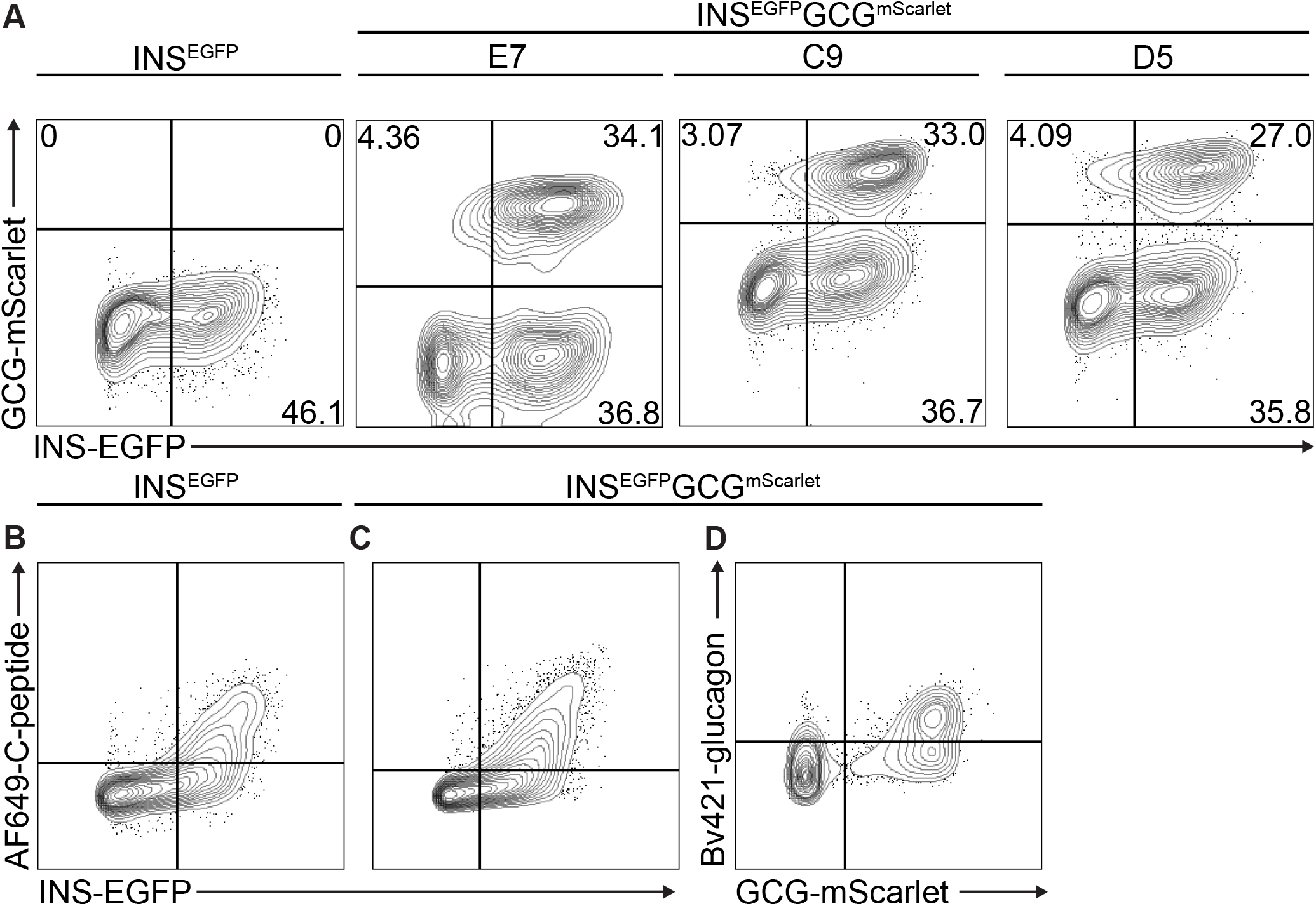
Flow cytometry analysis of INS^EGFP^ and three INS^EGFP^GCG^mScarlet^ clones. (A) Representative flow cytometry plots showing the differentiation efficiencies (% of alive cells) of the INS^EGFP^ and INS^EGFP^GCG^mScarlet^ Clones E7,C9 and D5 at the maturing endocrine cell stage (Stage 6 Day 21) (B,C) Representative flow cytometry plots illustrating the INS-EGFP reporter efficiency of maturing endocrine cells (Stage 6 Day 21) derived from (B) INS^EGFP^ and (C) INS^EGFP^GCG^mScarlet^ Clone E7 reporter cell lines. (D) Representative flow cytometry plot illustrating the GCG-mScarlet reporter efficiency of maturing endocrine cells (Stage 6 Day 21) derived from the INS^EGFP^GCG^mScarlet^ clone E7 reporter cell line.

## Supplementary Tables

**Table S1.**
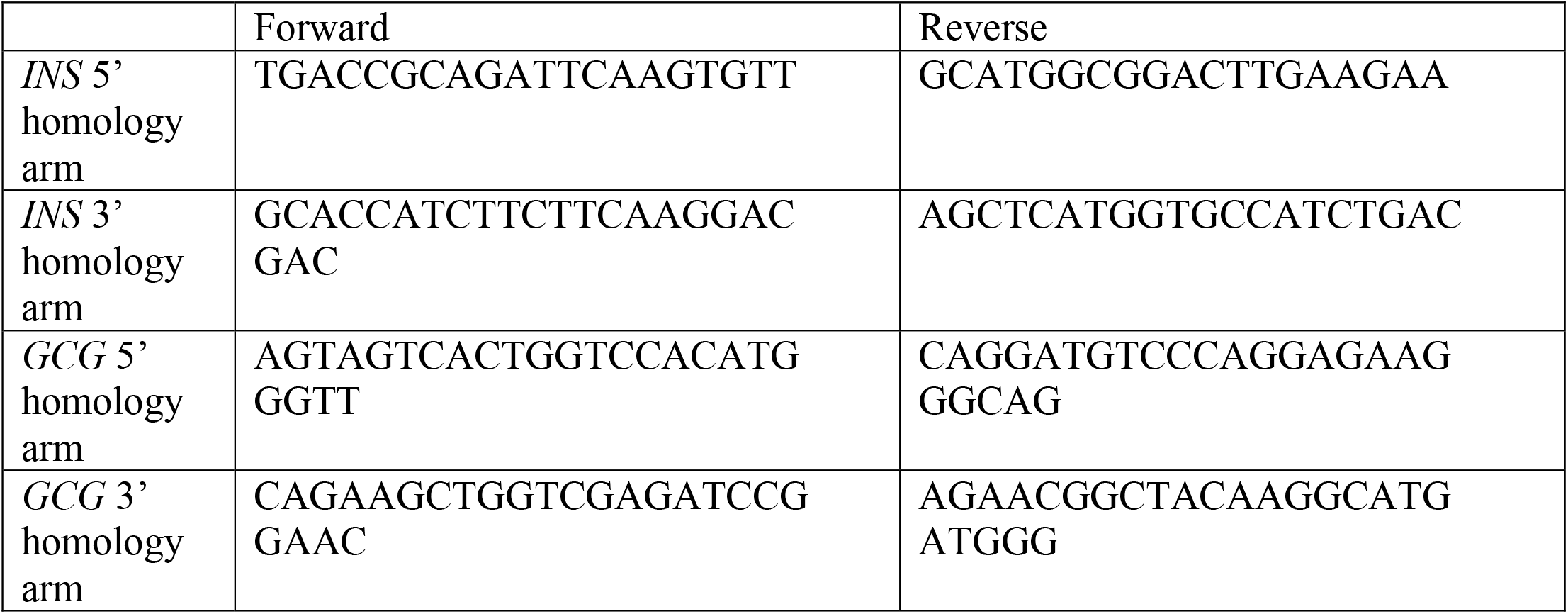
List of genotyping primers used.

**Table S2.**
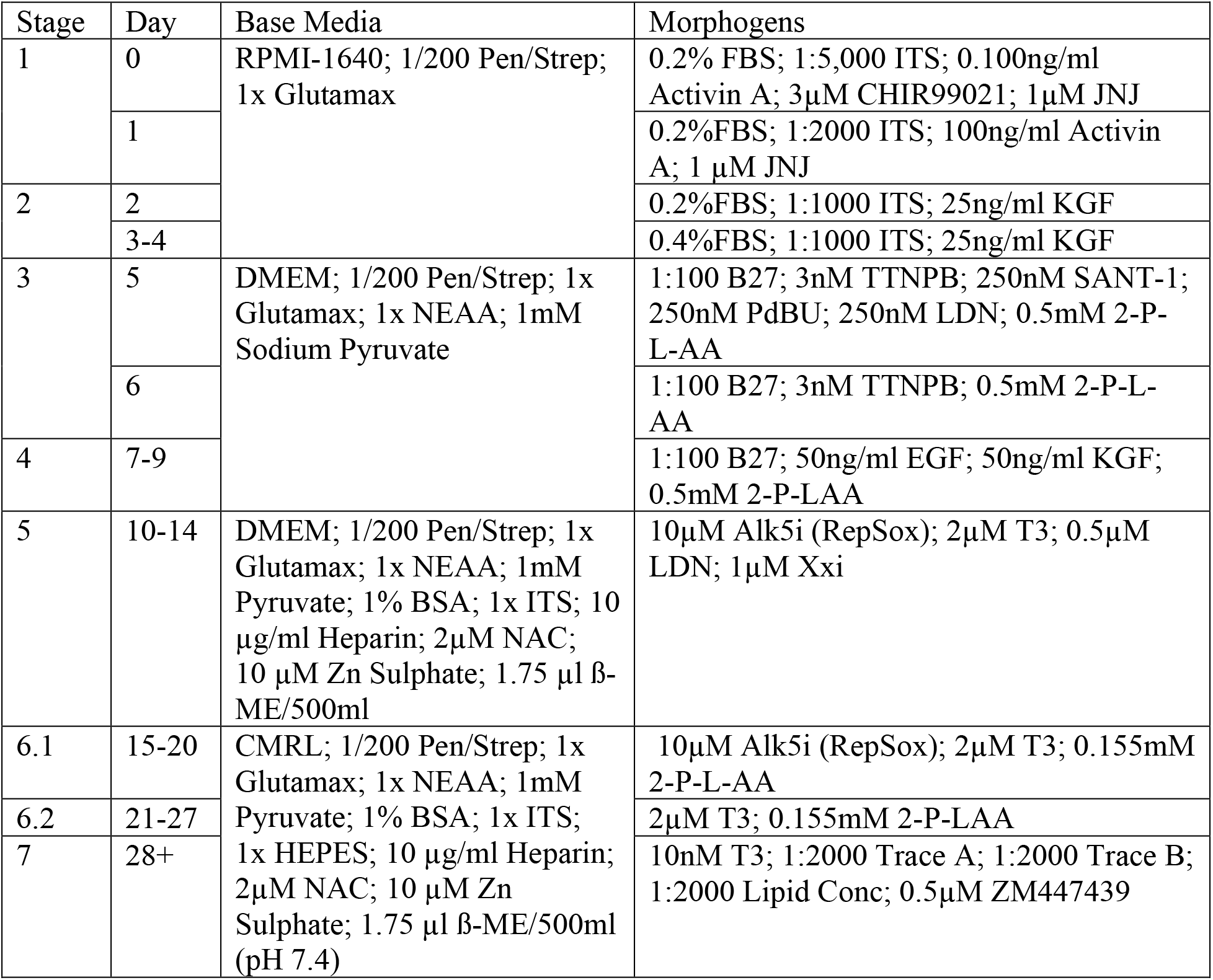
Media formulations.

**Table S3.**
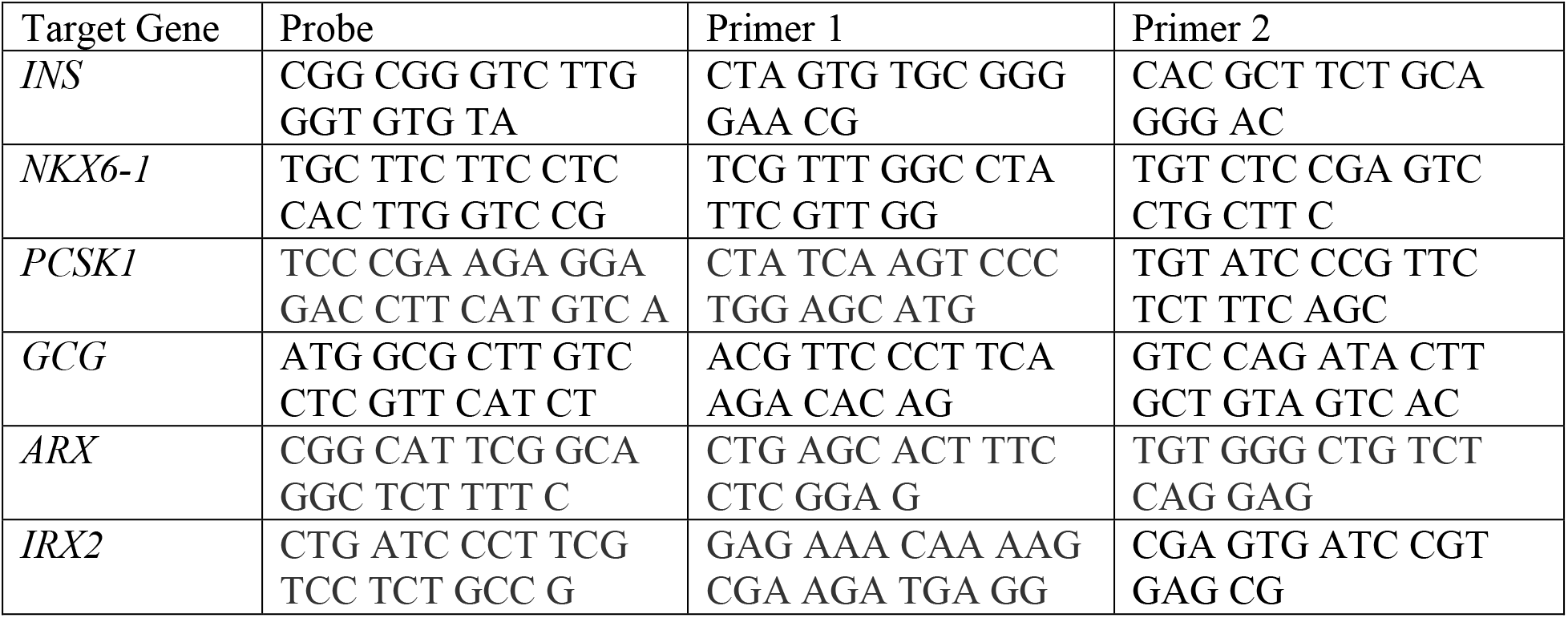
List of TAQMAN primers used.

**Table S4.**
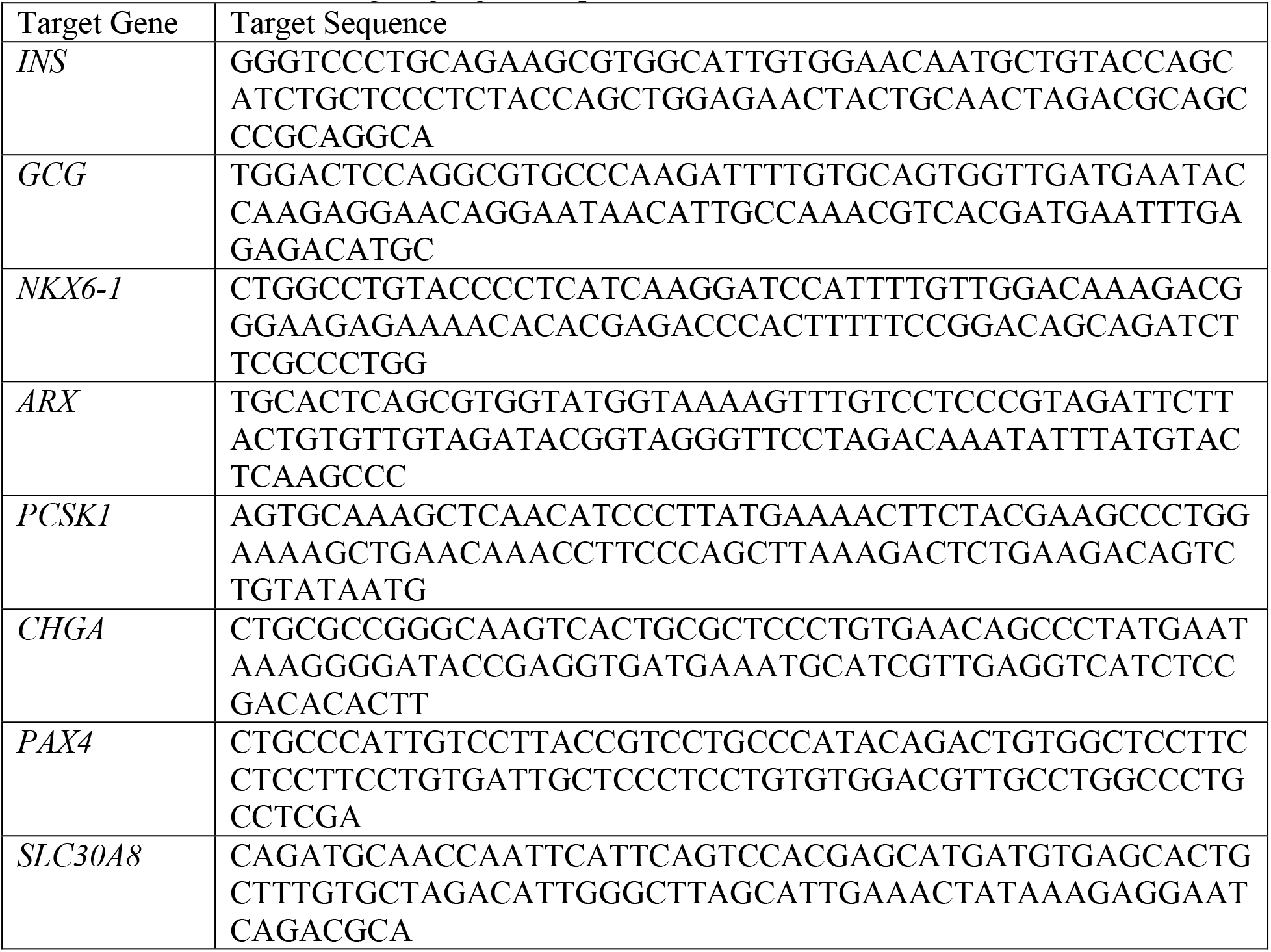
List of NanoString target gene sequences.

## References

Adams, M. T., & Blum, B. (n.d.). Determinants and dynamics of pancreatic islet architecture. Islets, 14(1), 82–100. https://doi.org/10.1080/19382014.2022.2030649

Alvarez-Dominguez, J. R., Donaghey, J., Rasouli, N., Kenty, J. H. R., Helman, A., Charlton, J., Straubhaar, J. R., Meissner, A., & Melton, D. A. (2020). Circadian Entrainment Triggers Maturation of Human In Vitro Islets. Cell Stem Cell, 26(1), 108–122.e10. https://doi.org/10.1016/j.stem.2019.11.011

Augsornworawat, P., Maxwell, K. G., Velazco-Cruz, L., & Millman, J. R. (2020). Single-Cell Transcriptome Profiling Reveals β Cell Maturation in Stem Cell-Derived Islets after Transplantation. Cell Reports, 32(8), 108067. https://doi.org/10.1016/j.celrep.2020.108067

Balboa, D., Barsby, T., Lithovius, V., Saarimäki-Vire, J., Omar-Hmeadi, M., Dyachok, O., Montaser, H., Lund, P.-E., Yang, M., Ibrahim, H., Näätänen, A., Chandra, V., Vihinen, H., Jokitalo, E., Kvist, J., Ustinov, J., Nieminen, A. I., Kuuluvainen, E., Hietakangas, V., … Otonkoski, T. (2022). Functional, metabolic and transcriptional maturation of human pancreatic islets derived from stem cells. Nature Biotechnology, 40(7), Article 7. https://doi.org/10.1038/s41587-022-01219-z

Basford, C. L., Prentice, K. J., Hardy, A. B., Sarangi, F., Micallef, S. J., Li, X., Guo, Q., Elefanty, A. G., Stanley, E. G., Keller, G., Allister, E. M., Nostro, M. C., & Wheeler, M. B. (2012). The functional and molecular characterisation of human embryonic stem cell-derived insulin-positive cells compared with adult pancreatic beta cells. Diabetologia, 55(2), 358–371. https://doi.org/10.1007/s00125-011-2335-x

Bellin, M. D., Barton, F. B., Heitman, A., Alejandro, R., & Hering, B. J. (2012). Potent induction immunotherapy promotes long-term insulin independence after islet transplantation in type 1 diabetes. American Journal of Transplantation, 12(6), 1576–1583. https://doi.org/10.1111/j.1600-6143.2011.03977.x

Blöchinger, A. K., Siehler, J., Wißmiller, K., Shahryari, A., Burtscher, I., & Lickert, H. (2020). Generation of an INSULIN-H2B-Cherry reporter human iPSC line. Stem Cell Research, 45, 101797. https://doi.org/10.1016/j.scr.2020.101797

Giudice, A., & Trounson, A. (2008). Genetic Modification of Human Embryonic Stem Cells for Derivation of Target Cells. Cell Stem Cell, 2(5), 422–433. https://doi.org/10.1016/j.stem.2008.04.003

Huang, L. T. (Helen), Zhang, D., Nian, C., & C, L. F. (2023). *Truncated CD19 as a selection marker for the isolation of stem cell derived β-cells* (p. 2023.04.05.535733). bioRxiv. https://doi.org/10.1101/2023.04.05.535733

Kelly, O. G., Chan, M. Y., Martinson, L. A., Kadoya, K., Ostertag, T. M., Ross, K. G., Richardson, M., Carpenter, M. K., D’Amour, K. A., Kroon, E., Moorman, M., Baetge, E. E., & Bang, A. G. (2011). Cell-surface markers for the isolation of pancreatic cell types derived from human embryonic stem cells. Nature Biotechnology, 29(8), Article 8. https://doi.org/10.1038/nbt.1931

Krentz, N. A. J., Nian, C., & Lynn, F. C. (2014). TALEN/CRISPR-Mediated eGFP Knock-In Add-On at the OCT4 Locus Does Not Impact Differentiation of Human Embryonic Stem Cells towards Endoderm. PLOS ONE, 9(12), e114275. https://doi.org/10.1371/journal.pone.0114275

Lavallard, V., Armanet, M., Parnaud, G., Meyer, J., Barbieux, C., Montanari, E., Meier, R., Morel, P., Berney, T., & Bosco, D. (2016). Cell rearrangement in transplanted human islets. The FASEB Journal, 30(2), 748–760. https://doi.org/10.1096/fj.15-273805

Liu, Z., Chen, O., Wall, J. B. J., Zheng, M., Zhou, Y., Wang, L., Ruth Vaseghi, H., Qian, L., & Liu, J. (2017). Systematic comparison of 2A peptides for cloning multi-genes in a polycistronic vector. Scientific Reports, 7(1), Article 1. https://doi.org/10.1038/s41598-017-02460-2

Micallef, S. J., Li, X., Schiesser, J. V., Hirst, C. E., Yu, Q. C., Lim, S. M., Nostro, M. C., Elliott, D. A., Sarangi, F., Harrison, L. C., Keller, G., Elefanty, A. G., & Stanley, E. G. (2012). INSGFP/whuman embryonic stem cells facilitate isolation of in vitro derived insulin-producing cells. Diabetologia, 55(3), 694–706. https://doi.org/10.1007/s00125-011-2379-y

Nair, G. G., Liu, J. S., Russ, H. A., Tran, S., Saxton, M. S., Chen, R., Juang, C., Li, M., Nguyen, V. Q., Giacometti, S., Puri, S., Xing, Y., Wang, Y., Szot, G. L., Oberholzer, J., Bhushan, A., & Hebrok, M. (2019). Recapitulating endocrine cell clustering in culture promotes maturation of human stem-cell-derived β cells. Nature Cell Biology, 21(2), 263–274. https://doi.org/10.1038/s41556-018-0271-4

Novakovsky, G., Sasaki, S., Fornes, O., Omur, M. E., Huang, H., Bayly, C. L., Zhang, D., Lim, N., Cherkasov, A., Pavlidis, P., Mostafavi, S., Lynn, F. C., & Wasserman, W. W. (2023). In silico discovery of small molecules for efficient stem cell differentiation into definitive endoderm. Stem Cell Reports, 18(3), 765–781. https://doi.org/10.1016/j.stemcr.2023.01.008

Peterson, Q. P., Veres, A., Chen, L., Slama, M. Q., Kenty, J. H. R., Hassoun, S., Brown, M. R., Dou, H., Duffy, C. D., Zhou, Q., Matveyenko, A. V., Tyrberg, B., Sörhede-Winzell, M., Rorsman, P., & Melton, D. A. (2020). A method for the generation of human stem cell-derived alpha cells. Nature Communications, 11(1), Article 1. https://doi.org/10.1038/s41467-020-16049-3

Rezania, A., Bruin, J. E., Xu, J., Narayan, K., Fox, J. K., O’Neil, J. J., & Kieffer, T. J. (2013). Enrichment of human embryonic stem cell-derived NKX6.1-expressing pancreatic progenitor cells accelerates the maturation of insulin-secreting cells in vivo. STEM CELLS, 31(11), 2432–2442. https://doi.org/10.1002/stem.1489

Rezania, A., Riedel, M. J., Wideman, R. D., Karanu, F., Ao, Z., Warnock, G. L., & Kieffer, T. J. (2011). Production of Functional Glucagon-Secreting α-Cells From Human Embryonic Stem Cells. Diabetes, 60(1), 239–247. https://doi.org/10.2337/db10-0573

Riedel, M. J., Asadi, A., Wang, R., Ao, Z., Warnock, G. L., & Kieffer, T. J. (2012). Immunohistochemical characterisation of cells co-producing insulin and glucagon in the developing human pancreas. Diabetologia, 55(2), 372–381. https://doi.org/10.1007/s00125-011-2344-9

Riopel, M., Li, J., Fellows, G. F., Goodyer, C. G., & Wang, R. (2014). Ultrastructural and immunohistochemical analysis of the 8-20 week human fetal pancreas. Islets, 6(4), e982949. https://doi.org/10.4161/19382014.2014.982949

Russ, H. A., Parent, A. V., Ringler, J. J., Hennings, T. G., Nair, G. G., Shveygert, M., Guo, T., Puri, S., Haataja, L., Cirulli, V., Blelloch, R., Szot, G. L., Arvan, P., & Hebrok, M. (2015). Controlled induction of human pancreatic progenitors produces functional beta-like cells in vitro. The EMBO Journal, 34(13), 1759–1772. https://doi.org/10.15252/embj.201591058

Sabatini, P. V., Speckmann, T., Nian, C., Glavas, M. M., Wong, C. K., Yoon, J. S., Kin, T., Shapiro, A. M. J., Gibson, W. T., Verchere, C. B., & Lynn, F. C. (2018). Neuronal PAS Domain Protein 4 Suppression of Oxygen Sensing Optimizes Metabolism during Excitation of Neuroendocrine Cells. Cell Reports, 22(1), 163–174. https://doi.org/10.1016/j.celrep.2017.12.033

Shapiro, A. M. J. (2012). Islet Transplantation in Type 1 Diabetes: Ongoing Challenges, Refined Procedures, and Long-Term Outcome. The Review of Diabetic Studies : RDS, 9(4), 385–406. https://doi.org/10.1900/RDS.2012.9.385

Shapiro, A. M. J., Pokrywczynska, M., & Ricordi, C. (2017). Clinical pancreatic islet transplantation. Nature Reviews Endocrinology, 13(5), Article 5. https://doi.org/10.1038/nrendo.2016.178

Shapiro, A. M. J., & Verhoeff, K. (2023). A spectacular year for islet and stem cell transplantation. Nature Reviews Endocrinology, 19(2), Article 2. https://doi.org/10.1038/s41574-022-00790-4

Sneddon, J. B., Tang, Q., Stock, P., Bluestone, J. A., Roy, S., Desai, T., & Hebrok, M. (2018). Stem Cell Therapies for Treating Diabetes: Progress and Remaining Challenges. Cell Stem Cell, 22(6), 810–823. https://doi.org/10.1016/j.stem.2018.05.016

Velazco-Cruz, L., Song, J., Maxwell, K. G., Goedegebuure, M. M., Augsornworawat, P., Hogrebe, N. J., & Millman, J. R. (2019). Acquisition of Dynamic Function in Human Stem Cell-Derived β Cells. Stem Cell Reports, 12(2), 351–365. https://doi.org/10.1016/j.stemcr.2018.12.012

Veres, A., Faust, A. L., Bushnell, H. L., Engquist, E. N., Kenty, J. H.-R., Harb, G., Poh, Y.-C., Sintov, E., Gürtler, M., Pagliuca, F. W., Peterson, Q. P., & Melton, D. A. (2019). Charting cellular identity during human in vitro β-cell differentiation. Nature, 569(7756), Article 7756. https://doi.org/10.1038/s41586-019-1168-5

